# A PAK kinase family member and the Hippo/Yorkie pathway modulate WNT signaling to functionally integrate body axes during regeneration

**DOI:** 10.1101/2022.07.14.500084

**Authors:** Viraj Doddihal, Frederick G. Mann, Eric Ross, Sean A. McKinney, Alejandro Sánchez Alvarado

**Affiliations:** Stowers Institute for Medical Research, Kansas City, MO, 64110, USA; Howard Hughes Medical Institute, Kansas City, MO, 64110, USA

**Keywords:** Body axes, regeneration, signaling, integration, WNT, Hippo

## Abstract

Successful regeneration of missing tissues requires seamless integration of positional information along the body axes. Planarians, which regenerate from almost any injury, use conserved, developmentally important signaling pathways to pattern the body axes. However, the molecular mechanisms which facilitate crosstalk between these signaling pathways to integrate positional information remain poorly understood. Here, we report a *p21-activated kinase* (*smed-pak1*) which functionally integrates the anterior-posterior (AP) and the medio-lateral (ML) axes. *pak1* inhibits WNT/β-catenin signaling along the AP axis and, functions synergistically with the β-catenin-independent WNT signaling of the ML axis. Furthermore, this functional integration is dependent on *warts* and *merlin* - the components of the Hippo/Yorkie (YKI) pathway. Hippo/YKI pathway is a critical regulator of body size in flies and mice, but our data suggest the pathway is required to pattern body axes in planarians. Our study provides a signaling network integrating positional information which can mediate coordinated growth and patterning during planarian regeneration.

## Introduction

The ability to restore lost tissues varies extensively among metazoans with some animals possessing little to no capacity to restore missing tissues, while others can regenerate complete animals from fragments removed from their bodies (Sánchez Alvarado and Tsonis, 2006; Sasidharan and Sánchez Alvarado, 2021). In regenerating animals, adult stem cells proliferate in response to injury and require positional cues that guide cell differentiation to replace missing tissues (Sánchez Alvarado and Tsonis, 2006). In axolotls, the nerve endings at the site of injury are sufficient to induce cell proliferation but require positional information to complete limb regeneration (Tanaka, 2016). Understanding the molecular basis of positional information in adult animals and how it is reset during regeneration is a fundamental and still unanswered question in regeneration biology. Multiple factors such as WNT, FGF, BMP, Sonic Hedgehog and Retinoic Acid, which provide positional information in embryonic tissues have similar roles in regeneration (Poss, 2010). However, the mechanisms by which the positional cues emanating from both wounded and uninjured tissues are coordinated and integrated to restore form and function remain unclear.

Planarian flatworms can regenerate complete animals from a fragment as small as 1/279^th^ the size of an animal (Morgan, 1898) or the equivalent of 8-10 thousand cells (Benham-Pyle et al., 2021). Such remarkable ability to regenerate is in part attributed to adult pluripotent planarian stem cells known as neoblasts (Bardeen and Baetjer, 1904; Dubois, 1949; Reddien et al., 2005). Upon amputation, neoblasts proliferate and differentiate to replace missing body parts based on positional information provided by muscle and other tissues (Bohr et al., 2021; Witchley et al., 2013). In planarians the anterior-posterior (AP) axis is patterned by the WNT/β-catenin, Hedgehog, and Activin signaling pathways (Adell et al., 2009; Cloutier et al., 2021; Gurley et al., 2008; Petersen and Reddien, 2008, 2009; Rink et al., 2009); while head and trunk identities along the AP axis are defied by FGFRL-mediated signaling (Cebrià et al., 2002; Lander and Petersen, 2016; Scimone et al., 2016). On the other hand, the dorsal-ventral (DV) axis of the animal is defined and maintained by BMP signaling, which is also essential for regeneration of the dorsal midline (Gaviño and Reddien, 2011; Molina et al., 2007; Reddien et al., 2007; Scimone et al., 2022). In addition to BMP signaling, the medio-lateral axis (ML) is regulated by β-catenin-independent WNT and Slit-mediated signaling (Adell et al., 2009; Almuedo-Castillo et al., 2011; Cebrià et al., 2007; Gurley et al., 2010). Altogether, these signaling pathways regulate the three body axes of the animal, and ultimately form the molecular coordinate system that sets up the planarian body plan.

Despite the great progress made in defining the pathways underpinning the maintenance and regeneration of adult body axes in planarians, the mechanisms by which these signals are processed and integrated remain poorly understood. For instance, planarians display the remarkable attribute of growing and degrowing depending on nutritional status. When fed, planarians increase their cell number and grow, but will shrink in size when starved all the while maintaining their scalar proportions throughout this process (Baguñà et al., 1990; Baguñá and Romero, 1981; Oviedo et al., 2003; Takeda et al., 2009; Thommen et al., 2019). In flies and mice, the Hippo/Yorkie pathway is the crucial regulator of cell number and organ size during both animal development and tissue homeostasis. The pathway consists of a kinase cascade involving Hippo (HPO), Salvador (SAV), Warts (WTS), and Merlin (MER) which inhibit Yorkie (YKI), the transcriptional cofactor that mediates the output of the pathway. Genetic inactivation of genes in the kinase cascade leads to hyperactivation of YKI, causing increased cell proliferation and decreased cell death leading to overgrowth of tissues and organs (Halder and Johnson, 2011; Harvey and Hariharan, 2012; Kango-Singh and Singh, 2009; Zhao et al., 2011). However, there is no known function for the Hippo/YKI pathway in regulating planarian body size. Instead, YKI is required for rescaling body axes during planarian regeneration (Lin and Pearson, 2014, 2017). Nevertheless, the process by which Hippo/YKI signaling may interact with the signaling pathways that pattern the planarian body axes remains unclear.

Most signal transduction pathways utilize protein phosphorylation as a dynamic modulator of functional output (Cohen, 2001; Olsen et al., 2006). For example, in the absence of WNT ligand, β-catenin is phosphorylated and is targeted for proteasomal degradation (Logan and Nusse, 2004), while binding of BMP ligand to its receptor results in phosphorylation of the transcriptional activator SMAD resulting in its nuclear localization and activation of target genes (Heldin et al., 1997). Protein phosphorylation and dephosphorylation are brought about by two classes of enzymes called kinases and phosphatases, respectively (Cohen, 2002; Hunter, 1995). A kinase or a phosphatase can target multiple proteins and alter their phosphorylation status, facilitating crosstalk between multiple signaling pathways (Cohen, 1992, 2002). In developing *Xenopus* embryos, Glycogen synthase kinase 3 (GSK3) phosphorylates β-catenin and Smad to inhibit both WNT and BMP signaling, thus modulating instructive signals for the establishment of the AP and the DV axes (Eivers et al., 2008; Fuentealba et al., 2007). As in embryogenesis, a regenerating planarian fragment needs to integrate a combination of signals to appropriately build lost tissues. Thus, we hypothesized that kinases and/or phosphatases may play key roles in coordinating the multiple signaling pathways driving the patterning of regenerating tissues.

We tested this hypothesis by performing an RNAi screen for kinases and phosphatases in the planarian flatworm *Schmidtea mediterranea* and identified that *p21 activated kinase 1 (pak1)* is necessary for proper patterning of both the AP and ML axes during regeneration. Our experiments revealed that *pak1* inhibits WNT/β-catenin signaling and is required for head regeneration at anterior blastema (Fig 1). In the ML axis, *pak1* synergizes with the β-catenin-independent WNT signaling to restrict the width of the midline. Furthermore, *pak1(RNAi)* phenotypes require *mer* and *wts,* which are components of the Hippo/YKI signaling pathway. Taken together, these results suggest that *pak1* patterns the AP and ML axes by facilitating crosstalk among the β-catenin-dependent and -independent WNT pathways and the Hippo/YKI pathway.

**Figure 1:**
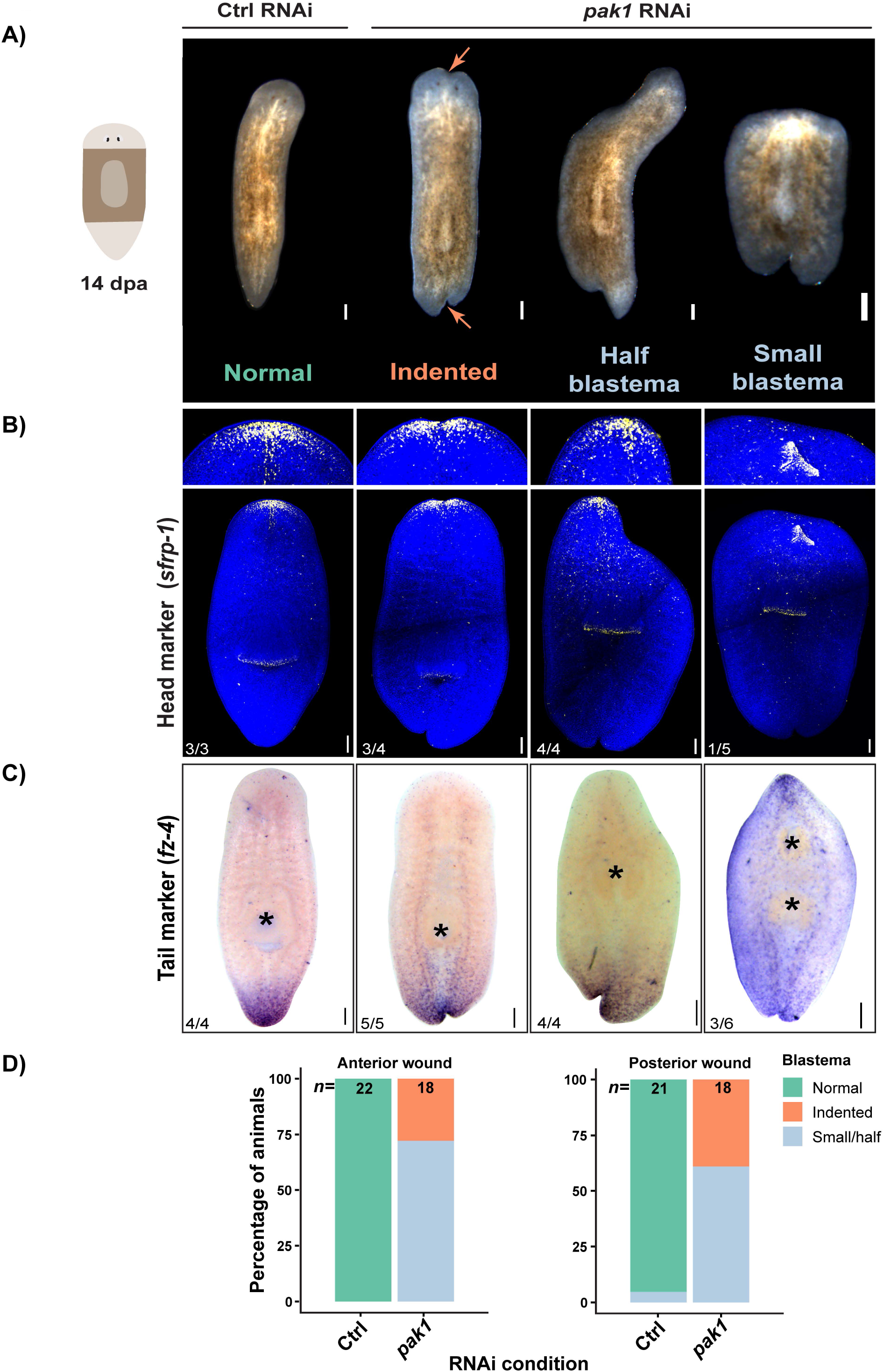
*pak1* is required for patterning the AP axis. **(A)** Live animal images of regenerating trunk fragments at 14 dpa. Orange arrows indicate midline indentations. **(B)** Maximum intensity projection images showing expression of head marker *sfrp-1.* **(C)** Expression of tail marker *fz-4.* Asterisk indicates pharynx. **(D)** Quantification of phenotype frequencies of both anterior and posterior blastema at 14 dpa. Scale bar: 200 um. See also Figures S1-S3.

## Results

### RNAi screen identified kinases and phosphatases important for blastema patterning

We utilized hidden Markov models of protein kinase and phosphatase domains to annotate the kinases and phosphatases in the planarian *S. mediterranea* (see experimental procedures). Using this method, we identified 619 putative kinases and 165 putative phosphatases (Table S1). Since the signals that set up and pattern the body axes are known to be expressed in differentiated tissues (Scimone et al., 2017; Witchley et al., 2013), we chose to screen only those genes whose expression could be detected in post-mitotic cells. To enrich for such genes, we looked for annotated kinases and phosphatases expressed in a lethally irradiated animal devoid of stem cells. All genes with a RPKM value ≥1 in a 4-day post irradiation bulk RNA-seq dataset (Cheng et al., 2018) were selected for a two-part RNAi screen (Fig S1A). Of those 604 genes which met the criteria, we cloned and screened 282 genes. For the screen, animals were fed with RNAi food 3 times, and scored after 14 days to identify genes that specifically manifested phenotypes in regenerating, but not unamputated animals (Fig S1B, S1C).

Out of the 282 genes screened, 7 genes were essential. RNAi of these genes caused lesions on dorsal epidermis or sometimes on the pharynx, leading to complete lysis by 14 days in homeostatic (unamputated) condition (Fig S1D, S1E, Table S1). Additionally, we observed RNAi phenotypes for 2 phosphatases and 4 kinases, which affected different aspects of regeneration but not homeostasis (Fig S1F, Table S1). Of the genes whose inhibition affected regeneration, two kinases, namely *Smed-tgfβr1* and *Smed-pak1* were specifically required for patterning the regenerating fragment and thus were chosen for further characterization.

### *tgfβr1* is required to pattern the dorsal-ventral axis

By 14 days post amputation (dpa) *tgfβr1(RNAi)* animals regenerated with midline indentations, tissue outgrowth on blastema, and ectopic photoreceptors (Fig S1F, S2A). Additionally, the dorsal epidermis of these animals displayed a ventral-like cilia pattern suggesting loss of DV polarity (Fig S2B). This prompted us to examine the role of *tgfβr1* in patterning the DV axis. The DV axis in planarians is defined and maintained by high BMP signaling on the dorsal side and, loss of BMP signaling leads to ventralization of the dorsal side resulting in ventral-like cilia pattern on dorsal epidermis, and formation of ectopic photoreceptors (Reddien et al., 2007). As *tgβr1(RNAi)* animals closely phenocopied *bmp(RNAi)* animals, we hypothesized that *tgfβr1* is required for BMP signaling. BMP signaling is restricted to the dorsal side by the expression of *admp* on the ventral side; loss of Bmp signaling leads to ectopic expression of *admp* (Gaviño and Reddien, 2011). Consistent with this model, loss of *tgfβr1* also resulted in ectopic expression of *admp* (Fig S2C) suggesting that *tgfβr1* kinase is essential for Bmp signaling. Given that TGFβR1 is predicted to be a receptor with transmembrane domains (Fig S2D), we propose that it functions as a receptor for BMP ligand and thus is critical for patterning of the DV axis of the animal.

### Blastema formation in *pak1(RNAi)* animals is delayed

*Smed-pak1* was the other gene discovered in the screen to be important for patterning of the regeneration blastema. Smed-PAK1 is a homolog of the <u>p</u>21-<u>a</u>ctivated <u>k</u>inase family of proteins, which possesses an N-terminal GTPase binding domain and a C-terminal kinase domain (Fig S3A, S3B) (Rane and Minden, 2014). Planarian *S. mediterranea* has at least 6 of *pak* kinase genes and we observed that one of them, *Smed-pak1* (SMESG000036933.1) is important for regeneration. Under normal conditions, amputation triggers migration of neoblasts to the wound site, which then proliferate to give rise to an unpigmented and undifferentiated structure called a blastema (Newmark and Sánchez Alvarado, 2002; Reddien and Sánchez Alvarado, 2004). The blastema becomes readily apparent by 3 dpa, and in due course differentiates and regenerates the missing tissues. However, amputation of *pak1(RNAi)* animals failed to form a detectable blastema even after 5 dpa (Fig S3C). Failure to form blastema can be a result of reduced stem cell proliferation, failed neoblast migration to the site of injury, disrupted neoblast differentiation or failed patterning (Elliott and Sánchez Alvarado, 2013; Scimone et al., 2014; Vásquez-Doorman and Petersen, 2014; Vogg et al., 2014). We measured neoblast proliferation in *pak1(RNAi)* animals using anti-phospho-Histone H3(Ser10) (H3P) antibody, which marks cells in the G2/M phase of cell cycle (Hendzel et al., 1997; Newmark and Sánchez Alvarado, 2000). The density of dividing cells in *pak1(RNAi)* animals was comparable to that of control animals, suggesting that delayed blastema formation was unlikely to be a consequence of reduced cell proliferation (Fig S3D). We also tracked the neoblast population during regeneration by fluorescent *in situ* hybridization (FISH) using the neoblast marker *piwi-1*. We failed to notice any qualitative differences in the number of neoblasts and observed that neoblasts were able to accumulate at the wound site by 1 dpa (Fig S3E). In addition to new tissue formation by cell proliferation, planarian regeneration involves remodeling of pre-existing tissues in a process called morphallaxis (Morgan, 1898; Reddien and Sánchez Alvarado, 2004). In a tail fragment that is regenerating anterior structures, the stem cell compartment is reorganized to make space for the regenerating pharynx. *pak1(RNAi)* fragments reorganized neoblasts to create a region devoid of stem cells even in the absence of blastema outgrowth (Fig S3E). However, by 9 dpa some *pak1(RNAi)* animals had grown small blastemas and formed photoreceptors suggesting successful differentiation (Fig S3C). Yet, the blastemas were irregularly shaped and had supernumerary photoreceptors. Hence, we hypothesized that *pak1* is likely required for patterning the regeneration blastema.

### *pak1* is necessary for head formation at anterior blastema

After 14 dpa, *pak1(RNAi)* animals exhibited a range of phenotypes affecting both anterior and posterior blastema. At lower doses of RNAi (3 feedings) we observed that most animals regenerated with indentations on the midline (Fig 1A). When subjected to higher doses of RNAi (6 feedings), in addition to indented blastema, animals exhibited either an asymmetric growth of their blastema which we termed “half blastema” or had small to no blastema (Fig 1A). Since a subset of RNAi animals had small blastema, we tested whether *pak1* was necessary for defining heads and tails. As in control animals, *pak1(RNAi)* animals with indented blastema and “half blastema” expressed the head marker *sfrp-1* at anterior ends (Fig 1B), and the tail marker *fz-4* at posterior ends (Fig 1C). However, animals with small blastema failed to express anterior *sfrp-1* and about half of them had *fz-4* expression at both anterior and posterior ends, indicating tail formation instead of head regeneration (Fig 1B, 1C). These blastema phenotype classes were quantified, and we observed that a majority of *pak1(RNAi)* animals showed either half or small blastema phenotypes (Fig 1D). A fraction of small blastema animals with two-tails formed inverted supernumerary pharynges anterior to pre-existing ones (5/14). A similar phenotype has been observed associated with reversal of polarity at anterior wounds in *apc(RNAi)* and *ptc(RNAi)* animal (Gurley et al., 2008; Rink et al., 2009). This strongly suggests that anterior wounds are likely taking on posterior identity in *pak1(RNAi)* animals. Consistent with loss of head fate at anterior ends, small/no blastema *pak1(RNAi)* animals failed to form cephalic ganglia and photoreceptors as visualized with Phospho-Ser/Thr antibody (Fig S4A). The ‘half blastema’ *pak1(RNAi)* animals had small cephalic ganglia and one photoreceptor while indented *pak1(RNAi)* animals had two cephalic ganglia and photoreceptors similarly to control animals (Fig S4A). Taken together, these results indicate that *pak1* is likely required for head formation at anterior wounds.

Body wall musculature has been implicated in providing positional information to pattern different tissues in the body (Sarkar et al., 2022; Scimone et al., 2017; Witchley et al., 2013). Even though the patterning along the AP axis was defective in *pak1(RNAi)* animals, we did not observe any gross abnormalities in body wall musculature as visualized by 6G10-2C7 antibody (Ross et al., 2015). However, in small/no blastema *pak1(RNAi)* animals, we observed formation of a second mouth around which the musculature was reorganized to accommodate the opening (Fig. S4B). These observations indicate that *pak1* regulates patterning of head at anterior wounds without disrupting the body wall musculature. Hence, we next tested whether *pak1* influences signaling pathways that define AP axis.

### *pak1* inhibits WNT/**β**-catenin signaling

The AP axis in planarians is established and maintained by WNT/β-catenin signaling (Gurley et al., 2008; Rink et al., 2009). WNT ligands and Frizzled receptors are expressed at the posterior of the animal and WNT inhibitors *notum* and *sfrp-1* are expressed at the anterior of the animal (Gurley et al., 2008; Petersen and Reddien, 2008, 2009, 2011; Rink et al., 2009) (Fig 2A). The WNT ligand *wnt1* is induced as part of a general wound response from 6-24 hours post amputation (hpa), which is then selectively suppressed at anterior wounds. By 2 dpa, *wnt1* expression coalesces at the caudal end to form the posterior pole (Petersen and Reddien, 2009).

**Figure 2:**
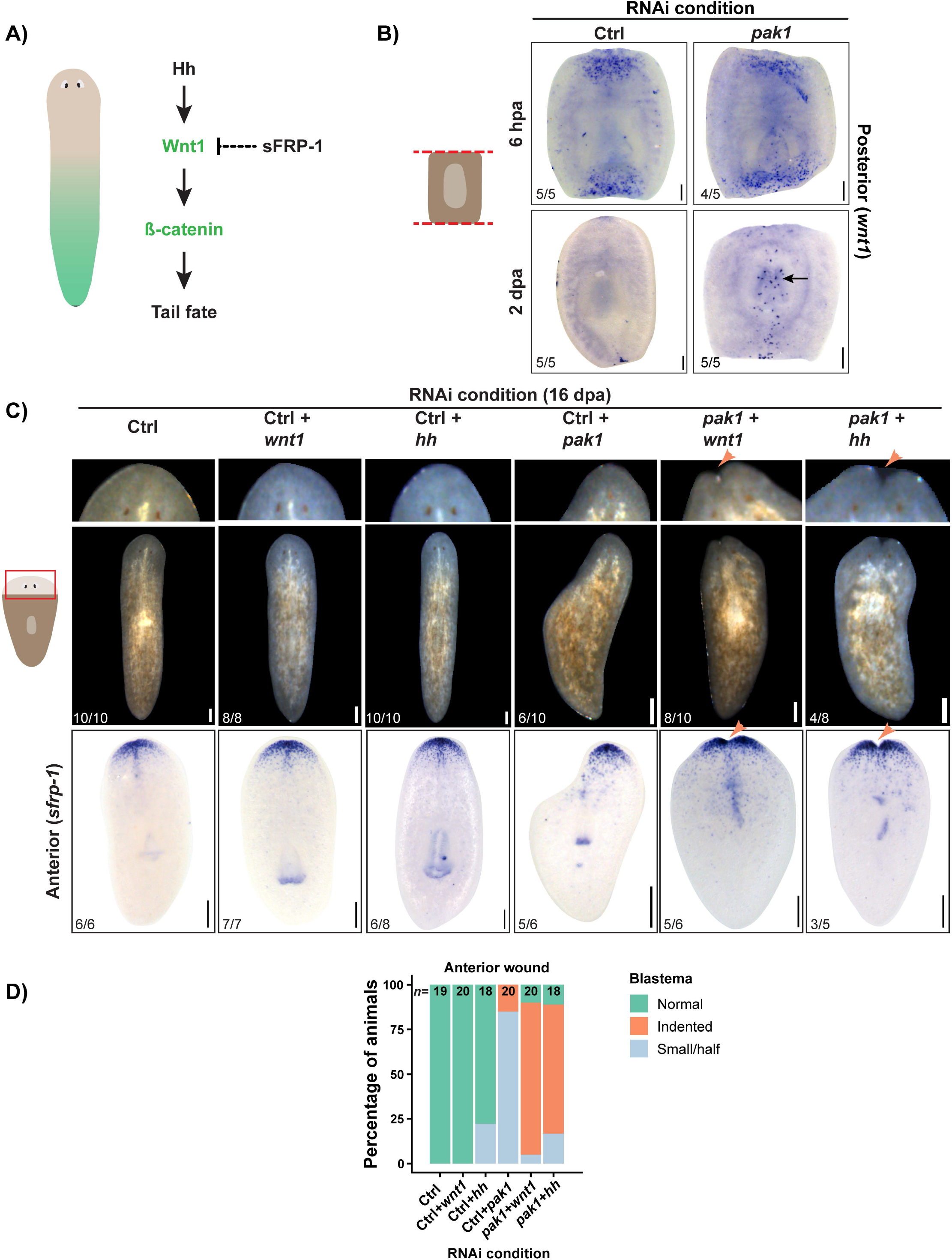
*pak1* inhibits the WNT/β-catenin signaling. **(A)** Graphical representation of the WNT/β-catenin gradient and the pathway defining the AP axis of the animal. **(B)** Expression of *wnt1* at 12 hpa and 2 dpa. Black arrow indicating ectopic expression of *wnt1*. **(C)** Live animal images of regenerated tail fragments at 16 dpa (top row) and anterior expression of *sfrp-1* (bottom row) from double RNAi experiment. Orange arrows indicate midline indentations. **(D)** Relative frequencies of different phenotypic classes of anterior blastema from 14 dpa trunk and tail fragments. Scale bar: 200 um. See also Figure S4.

Since *pak1(RNAi)* causes tail formation at anterior blastemas, we expected these animals display hyperactivated WNT/β-catenin signaling. To determine whether *pak1* affects the activation of *wnt1* after wounding or the suppression of *wnt1* at posterior poles, we performed *in situ* hybridization for *wnt1*. At 12 hpa *wnt1* expression was qualitatively similar in both control and RNAi animals, but at 2 dpa *pak1(RNAi)* animals had ectopic *wnt1* expression all along the AP axis of the regenerating fragment (Figure 2B). These results indicate that *pak1(RNAi)* does not affect the wound-induced WNT response but specifically causes hyperactivation of the WNT/β-catenin signaling patterning the AP axis.

If hyperactivation of *wnt1* were the reason for formation of tails at anterior blastemas, then reducing WNT/β-catenin signaling should rescue anterior head formation. We tested this hypothesis by performing double RNAi experiments wherein WNT/β-catenin signaling was reduced by knocking down either *hh, wnt1* or β*-catenin* in combination with *pak1(RNAi)*. These double RNAi conditions rescued anterior head regeneration, as confirmed by the restoration of *sfrp-1* expression at the anterior end (Fig 2C, 2D, S4C). These results support the hypothesis that *pak1* inhibits WNT/β-catenin signaling and allows for head regeneration at anterior blastema in planarians.

However, reduction of WNT/β-catenin signaling by introduction of either *hh, wnt1,* or β*-catenin* RNAi, were unable to rescue deformities of the posterior blastema generated by *pak1(RNAi)* (Fig S4E). Interestingly, most of the regenerated heads in these double RNAi conditions had an indentation on the midline (Fig 2D). This led us to explore the functions of *pak1* in patterning the medio-lateral (ML) axis.

### *pak1* inhibits lateral expansion of medial tissues

To test if the midline indentations were caused by a mispatterned ML axis, we performed FISH for the midline marker *slit* (Cebrià et al., 2007). In *hh(RNAi)* and *wnt1(RNAi)* animals, ventral expression of *slit* was indistinguishable from control animals. However, in *pak1(RNAi); hh(RNAi)* and *pak1(RNAi); wnt1(RNAi)* animals ventral *slit* expression was widened and failed to taper at the tips suggesting an expanded midline. Since RNAi of *wnt1* or *hh* alone did not result in widened midline, this indicates a possible role for *pak1* in patterning the ML axis (Fig S5A).

In *pak1(RNAi)* a subset of animals regenerated with an indented blastema (Fig 1A). In these animals, ventral *slit* expression was widened and failed to narrow down at the tips (Fig 3A), and the animals were wider when compared to control animals (Fig 3B). In planarians, the two cephalic ganglia are on either side of the *slit* expression domain and as *slit* expression tapers towards the anterior end, the two ganglia meet to form the anterior neural commissure (Fig 3C) (Cebrià et al., 2007; Gurley et al., 2010). Two photoreceptors are dorsally located and send their neural projections to the ganglia while forming an optic chiasma (Fig 3D). Indented *pak1(RNAi)* animals failed to form the anterior neural commissure and formed supernumerary photoreceptors which failed to make appropriate neural projections (Fig 3C, 3D). These phenotypes recapitulated the widened midline phenotypes previously observed in *wnt5(RNAi)* animals (Adell et al., 2009; Almuedo-Castillo et al., 2011; Gurley et al., 2010) indicating a role for *pak1* in regulating the ML axis in planarians.

**Figure 3:**
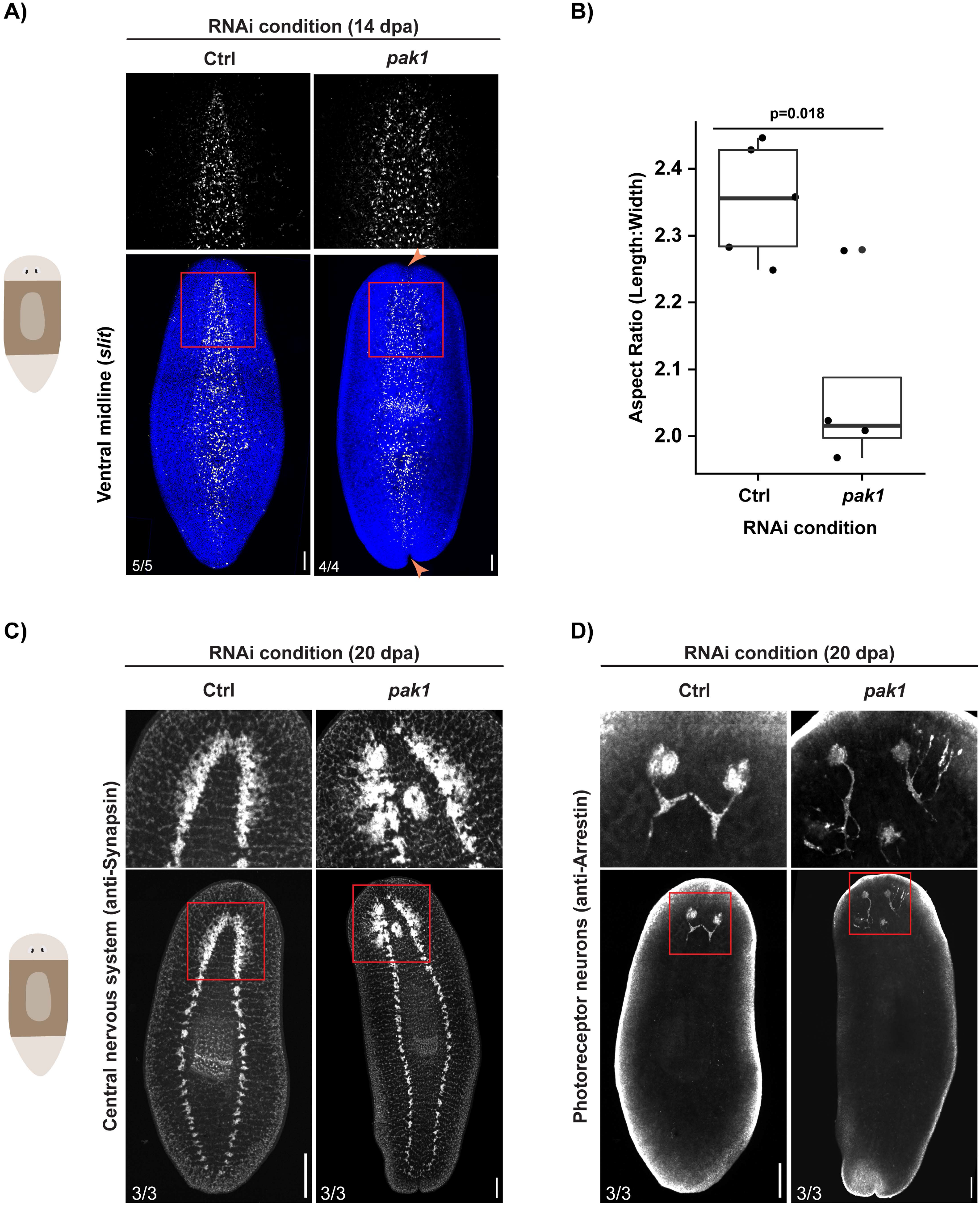
*pak1* regulates patterning of the ML axis. **(A)** Maximum intensity projection of ventral third of the animal showing midline (*slit)* in regenerated animals. Area marked by red box is shown in the top row. **(B)** Ratio of length and width measured from DAPI stained animals at 14 dpa. p-value is from two-tailed Student’s t-Test. **(C)** Central nervous system in 20 dpa animals. **(D)** Photoreceptors and optic chiasma in 20 dpa animals. Zoomed images (top row) are from the regions indicated by red boxes in the whole animal images (bottom row). Images in both (C) and (D) are maximum intensity projections. Scale bar: 200 um. See also Figure S5.

### *pak1* synergizes with **β**-catenin-independent WNT signaling

The ML axis in planarians is regulated by the midline expression of *slit* and lateral expression of the WNT ligand of the β-catenin-independent pathway, *wnt5*. *slit* and *wnt5* are mutually inhibitory, where *slit(RNAi)* results in expansion of *wnt5* expression towards the midline, and *wnt5(RNAi)* causes expansion of *slit* expression towards lateral edges (Fig 4A) (Adell et al., 2009; Gurley et al., 2010). Since *pak1(RNAi)* resulted in widened *slit* expression, we tested whether *pak1* functions along with *wnt5* to inhibit medial fates. In uninjured animals after 6 feedings of RNAi, neither *pak1(RNAi)* nor *wnt5(RNAi)* animals formed any ectopic photoreceptors, but *pak1(RNAi); wnt5(RNAi)* animals formed supernumerary photoreceptors (Fig 4B). Since double RNAi of *pak1* and *wnt5* resulted in a more severe phenotype compared to single RNAi, we concluded that both *pak1* and *wnt5* are synergistically functioning to inhibit lateral spread of medial tissues. Similar effects were obtained during regeneration, where a greater number of *pak1(RNAi); wnt5(RNAi)* animals regenerated with indented blastema when compared to *pak1(RNAi)* or *wnt5(RNAi)* animals (Fig S5B).

**Figure 4:**
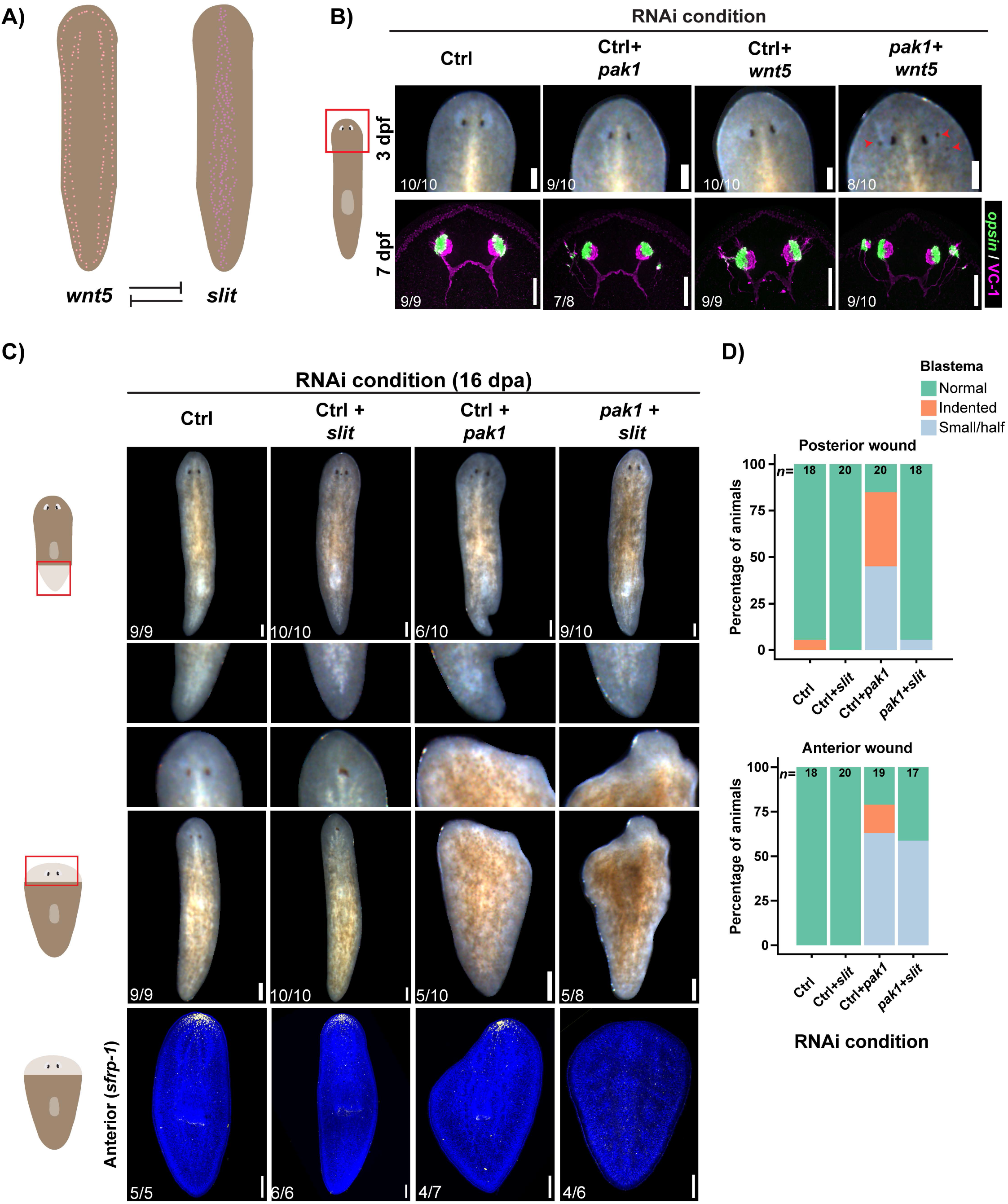
*pak1* synergizes with the β-catenin-independent WNT signaling. **(A)** Representation of expression patterns of genes that regulate ML axis. **(B)** Live animal images of uninjured animals at 3 dpf. Red arrowheads point to ectopic pigment cups (top row). Maximum intensity projection images showing photoreceptor neurons in 7 dpf uninjured animals (bottom row). **(C)** Live animal images with posterior regeneration (top two rows) and anterior regeneration (middle two rows) at 16 dpa. Expression of the anterior marker *sfrp-1* in the double RNAi conditions (bottom row). **(D)** Phenotypic frequencies of posterior and anterior blastema at 14 dpa. All fluorescent images are maximum intensity projections. Scale bar: 200 um. See also Figure S5.

Next, we tested whether the indented blastema phenotypes in *pak1(RNAi)* animals were caused by the widened *slit* expression domain. *pak1(RNAi); slit(RNAi)* rescued all indented blastema phenotypes at both anterior and posterior wounds (Fig 4C, 4D). However, *pak1(RNAi); slit(RNAi)* could not rescue anterior head regeneration suggesting failed head regeneration is not likely due to changes along the ML axis (Fig 4C, 4D). These results confirm that *pak1* is likely synergizing with the β-catenin-independent WNT signaling to shape the ML axis in planarians. Taken together we conclude that *pak1* is acting on both the β-catenin-dependent and -independent WNT pathways to shape the AP and ML axes respectively.

To explore mechanisms by which *pak1* is patterning the regeneration blastema we looked for genes which produced similar RNAi phenotypes. Interestingly, *pak1(RNAi)* phenocopied the previously reported *yki* loss-of-function phenotype, especially with the expanded expression of *wnt1* along the midline (Fig 5A) (Lin and Pearson, 2014, 2017). In mammals PAK1 is a known upstream activator of YAP (vertebrate homolog of YKI) (Sabra et al., 2017). Thus, we hypothesized that *pak1* may be regulating Hippo/YKI signaling to shape the body axes during planarian regeneration.

**Figure 5:**
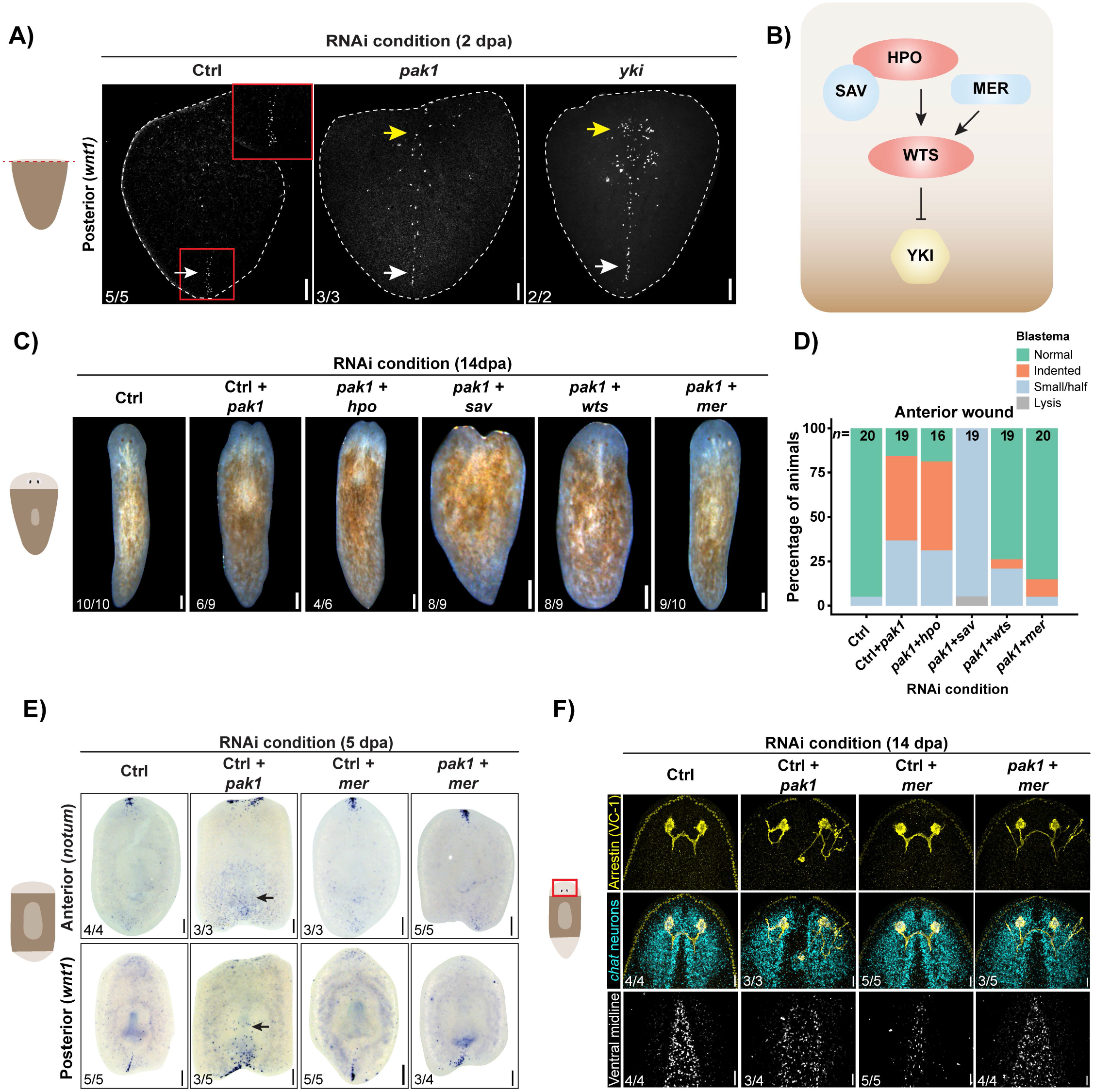
*pak1* functions with components of the Hippo/YKI signaling to shape AP and ML axis. **(A)** Expression of posterior determinant *wnt1* at 2 dpa. White arrow indicates WT expression and yellow arrow indicates ectopic expression. **(B)** Overview of Hippo/YKI signaling (adapted from (Yin et al., 2013)). **(C)** Anterior regeneration phenotypes of double RNAi animals in 14 dpa tail fragments. **(D)** Quantification of phenotypic classes of anterior blastema in double RNAi animals at 14 dpa. **(E)** Expression of anterior *notum* (top row) and posterior *wnt1* (bottom row) in 5 dpa trunk fragments. Black arrows indicate ectopic expression. **(F)** Head regions of trunk fragments at 14 dpa immunostained with anti-arrestin (VC-1) marking photoreceptor neurons and optic chiasma (yellow) and probed with *chat* to visualize cephalic ganglia (cyan). Ventral midline *slit* expression in trunk fragments at 14 dpa. Images in (A) and (F) are maximum intensity projections. Scale bar: 200 um. See also Figures S6.

### Components of the Hippo signaling pathway are required for *pak1(RNAi)* phenotype

HPO in association with SAV and MER phosphorylates and activates WTS, which phosphorylates YKI and inhibits expression of its target genes (Fig 5B) (Halder and Johnson, 2011; Yin et al., 2013; Yu and Guan, 2013). In mammals PAK1 can directly phosphorylate and inhibit MER, thus activating YAP/YKI (Rong et al., 2004; Sabra et al., 2017; Xiao et al., 2002). To test possible genetic interactions between *Smed-pak1* and components of the Hippo pathway, we performed epistasis experiments. RNAi of the members of the Hippo pathway alone did not affect regeneration, except for *sav(RNAi),* which affected blastema formation or caused lysis (Fig S6A). Rarely, knockdown of *wts* or *mer* resulted in tiny outgrowths at the tip of regenerating heads (Fig S6B). When testing the genetic interaction between components of the Hippo pathway and *pak1,* we did not observe rescue of posterior regeneration (Fig S6C). However, RNAi of the genes *wts* and *mer,* suppressed anterior phenotypes in *pak1(RNAi)* animals as most of these double RNAi animals regenerated heads without any midline indentations (Figure 5C, 5D). Thus, we suspect that *pak1* may inhibit *mer* and *wts* in planarians to pattern anterior blastema during regeneration.

### *pak1* and *mer* function to shape both AP and ML axes during regeneration

Since MER is a potential substrate for PAK1, we chose to further characterize *pak1(RNAi); mer(RNAi)* phenotype. As *pak1(RNAi)* animals failed to form anterior tissues, we also observed that these animals failed to regenerate the prepharyngeal population of secretory cells marked by *mag1*. This population of secretory cells did not regenerate in *pak1(RNAi); mer(RNAi)* animals (Fig S6D). Additionally, *pak1(RNAi)* animals had ectopic axonal projections, which persisted in *pak1(RNAi); mer(RNAi)* animals. These results indicate that *pak1* likely regulates photoreceptor axonal guidance and regeneration of prepharyngeal secretory cells independent of *mer* (Fig S6B). However, patterning along the AP and ML axes by *pak1* relied on *mer. pak1(RNAi)* animals failed to form an anterior pole expressing the WNT inhibitor *notum* while ectopically expanding *wnt1* and *notum* expression. We found that this phenotype was dependent on *mer* as *pak1(RNAi); mer(RNAi)* animals restored polarized expression of *wnt1* and *notum* (Figure 5E). For those RNAi animals which formed heads at the anterior plane of amputation, we computed ratios of tail to body lengths and found that *pak1(RNAi)* animals formed proportionally longer tails in a *mer* dependent manner (Fig S6E). These results collectively show that *mer* is required for elevated WNT/β-catenin signaling in *pak1(RNAi)* animals. Along the ML axis of the animal, knockdown of *pak1* results in widening of the ventral midline as visualized by *slit* expression. This results in the loss of the anterior neural commissure and optic chiasma. These phenotypes were rescued in *pak1(RNAi); mer(RNAi)* animals (Figure 5F). From these data, we conclude that *mer* is necessary for expansion of the midline in *pak1(RNAi)* animals.

Given that PAK1 is known to phosphorylate and inhibit MER in mammalian cells (Rong et al., 2004), we posit that *Smed-pak1* inhibits *Smed-mer* and subsequently activates *Smed-yki,* to restrict WNT/β-catenin signaling along the AP axis and to synergize with β-catenin-independent WNT signaling to shape the ML axis (Fig 6A, 6B). Taken together, our study has led us to propose a model where the AP and ML axes can be potentially interlinked by *pak1* and Hippo/YKI signaling, providing a possible mechanism for coordinated patterning and growth during animal regeneration (Fig 6C).

**Figure 6:**
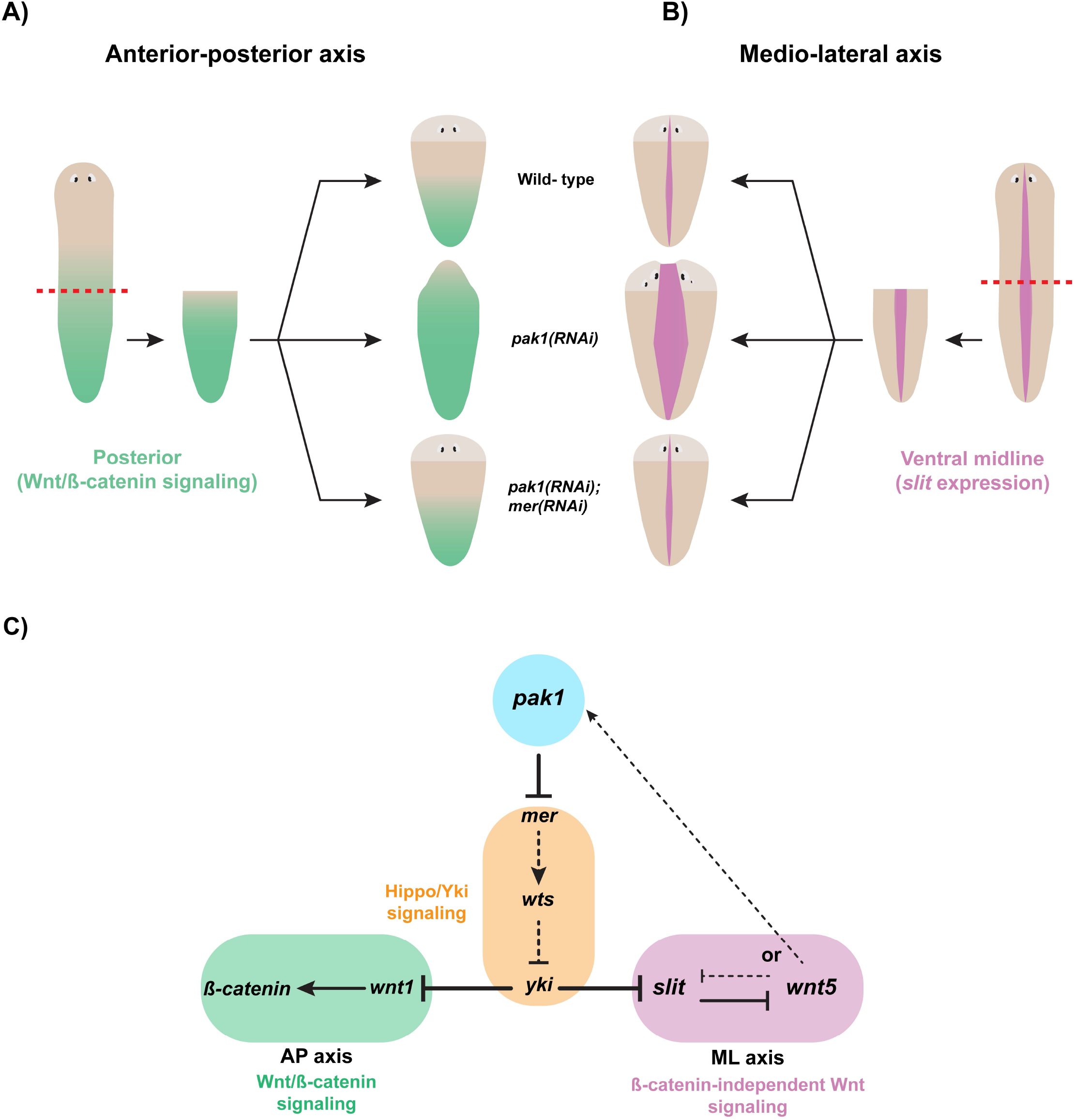
Integration of AP and ML axes by *pak1* and *mer*. **(A)** Cartoon describing the function of *pak1* and *mer* in patterning the WNT/β-catenin signaling along the AP axis. **(B)** Graphical representation of role of *pak1* and *mer* in shaping the ML axis by regulating ventral midline expression of *slit*. **(C)** Proposed signaling network of *pak1* which facilitates patterning of both the AP and ML axes by regulating the activity of Hippo/YKI, WNT/β-catenin, and β-catenin-independent WNT signaling pathways.

## Discussion

### Functional integration of the AP and ML axes

During regeneration, cells interpret a combination of signals emanating from both injured and uninjured tissues to restore the missing body parts. These signals not only instruct cell proliferation, but also provide positional cues so that regenerated tissues maintain polarity and scale in accordance with the rest of the body (Newmark and Sánchez Alvarado, 2002; Sánchez Alvarado and Tsonis, 2006). Positional cues from different body axes must be integrated for the coordinated growth and patterning of tissues (De Robertis, 2010). In this study, we showed that *pak1* functions with Hippo/YKI signaling components, *mer* and *wts,* to inhibit the WNT/β-catenin signaling of the AP axis and synergizes with the β-catenin-independent WNT signaling patterning the ML axis. Because systemic abrogation of *pak1* leads to patterning defects along both the AP and ML axes, we propose that *pak1* may be a ‘functional integrator’ of the pathways patterning these two orthogonal body axes.

One of the predictions of the model is that the β-catenin-independent WNT signaling of the ML axis inhibits the WNT/ β-catenin signaling of the AP axis via *pak1* and Hippo/YKI pathway (Fig 6C). Such an inhibition of the WNT/β-catenin signaling by β-catenin-independent WNT signaling has been previously observed in mammalian cells *in vitro,* during vertebrate embryogenesis, and hematopoietic stem cell maintenance (Mikels and Nusse, 2006; Park et al., 2015; Sugimura et al., 2012; Topol et al., 2003; Torres et al., 1996; Weidinger and Moon, 2003; Westfall et al., 2003). This potential signaling can provide an elegant mechanism in which the AP and ML axes are interlinked by *pak1* and Hippo/YKI pathway, allowing dynamic reestablishment and proportionate reshaping of body axes during regeneration. Future research will aim to elucidate the biochemical mechanism by which such integration may be effected at the cellular level.

### Regulation of body size by *pak1* and Hippo/YKI signaling

During planarian growth there is a net increase in cell number due to increased cell proliferation and minimized cell death. The opposite occurs during de-growth, where more cells are lost which results in shrinkage (Baguñà et al., 1990; Baguñá and Romero, 1981; Oviedo et al., 2003; Pellettieri et al., 2010; Takeda et al., 2009; Thommen et al., 2019). This balance between cell proliferation and cell death is maintained at least in part by insulin signaling, mTOR pathway and JNK signaling (Almuedo-Castillo et al., 2014; Miller and Newmark, 2012; Tu et al., 2012). In other organisms these pathways are known to function upstream of YKI/YAP, which is critical for cell number and organ size (Codelia et al., 2014; Csibi and Blenis, 2012; Ibar and Irvine, 2020; Sayedyahossein et al., 2020). However, in animals with indeterminate growth like planaria and *Hydra*, YKI/YAP patterns the body axes by modulating WNT signaling (Brooun et al., 2022; Lin and Pearson, 2014, 2017; Unni et al., 2021). In *Hydra*, WNT signaling indirectly regulates body size by patterning the oral-aboral axis (Mortzfeld et al., 2019). Thus, during indeterminate growth and de-growth, it is possible that YKI/YAP mediated scale regulation of body axes may play a role in determining organ size.

Apart from *smed*-*yki*, there are no reported roles for other members of the Hippo/YKI pathway in patterning of body axes in planarians. Our data indicate that in *pak1(RNAi)* background *wts* and *mer* affect patterning of the body axes (Fig 5B). We also observe that hyperactivation of WNT/β-catenin signaling and widening of midline *slit* expression in *pak1(RNAi)* is dependent on *mer* (Fig 5C). In fact, another potential modulator of Hippo/YKI signaling, *mob4* of the STRIPAK complex, is required to scale tail with respect to the size of body (Schad and Petersen, 2020). Taken together, we propose that *pak1* and Hippo/YKI signaling may be regulating organ size during growth and de-growth by modulating body axes.

### Mechanotransduction during regeneration

Besides shaping body axes, *Smed-pak1* is required for blastema formation. It is unclear whether this phenotype in *pak1(RNAi)* animals is due to anomalies along the body axes, considering that failure to form anterior or posterior poles is known to result in small blastemas (Scimone et al., 2014; Vásquez-Doorman and Petersen, 2014; Vogg et al., 2014). However, delay in blastema formation could also be due to impaired wound healing, as a similar process called ‘dorsal closure’ during *Drosophila* embryogenesis requires Pak1 (Conder et al., 2004; Harden et al., 1996). Furthermore, PAK kinases are known to transduce mechanical cues from ECM/Integrin interactions and facilitate body elongation (Labouesse, 2011; del Pozo et al., 2000; Zhang et al., 2011). In planarians, inhibition of Integrin signaling by β*1-integrin(RNAi)* also results in small blastema (Bonar and Petersen, 2017; Seebeck et al., 2017). Additionally, like β*1-integrin(RNAi)* animals, *pak1(RNAi)* animals develop ectopic neural ‘spheroids’ (Fig 3C), strongly suggesting these two genes function in the same process. Integrins and PAK kinases are known to transduce mechanical signals from ECM and regulate the activity of YKI/YAP (Chakraborty et al., 2017; Dupont et al., 2011; Sabra et al., 2017). With the development of new technologies to characterize planarian ECM, it is now feasible to identify upstream members of the ECM that are critical for mechanotransduction during planarian regeneration (Benham-Pyle et al., 2022; Sonpho et al., 2021).

### Pleiotropic functions of *pak1* and *mer*

Merlin is a cytoskeletal adaptor protein that can respond to mechanical cues and modulate Hippo/YKI signaling (Das et al., 2015; Yin et al., 2013). PAK1 can directly phosphorylate and inhibit Merlin or can regulate actin cytoskeleton which indirectly modulates the function of Merlin (Eby et al., 1998; Rane and Minden, 2014; Sabra et al., 2017; Yin et al., 2013). Apart from Hippo/YKI signaling, PAK1 and Merlin regulate other signaling pathways, which could be the reason for mispatterned posterior blastema or ectopic axonal projections in *pak1(RNAi); mer(RNAi)* animals (Fig S8C, S9B) (Bashaw and Klein, 2010; Kim et al., 2016; Lavado et al., 2014; Mota and Shevde, 2020). PAK1 and Merlin are extensively studied in the context of cancer, where PAK1 is activated in cancer cells and Merlin (NF2 – the vertebrate homolog) functions as a tumor suppressor (LaJeunesse et al., 1998; Rouleau et al., 1993; Trofatter et al., 1993; Ye and Field, 2012). It would be interesting to study the functions of these genes in maintenance and proliferation of adult stem cells in planarians. Future experiments identifying targets of Smed-PAK1, and other regulators of Smed-MER will help uncover the pleiotropic functions of these proteins.

## Conclusion

In this study, we identified a kinase that functionally integrates signals patterning the AP and ML axes. We demonstrated that *pak1* works with the components of the Hippo/YKI pathway to modulate β-catenin-dependent and -independent WNT signaling. This also revealed a potential linkage between the AP and ML axes which provide a mechanism for coordinated growth and regeneration. Furthermore, the Hippo/YKI pathway known to regulate organ size during embryonic development in flies and mice is likely functioning to pattern the body axes in adult planarians. Thus, in these adult animals with indeterminate growth and de-growth, the scalar proportion of body is maintained likely by shaping the body axes.

## Acknowledgements

We thank all the members of Sánchez Alvarado lab for regular discussions and advice. Thanks to Zainab Afzal, Alice Accorsi, Stephanie Nowotarski, and Manon Valet for comments on the manuscript. We also thank Cindy Maddera for help with imaging, all the members of Reptiles and Aquatics core for their efforts in culturing planarians and Ariel Bazzini, Matt Gibson, and Kausik Si for their insightful suggestions as part of V.D.’s thesis committee. A.S.A. is an investigator of the Howard Hughes Medical Institute (HHMI) and the Stowers Institute for Medical Research. This work was supported by (funding information) to A.S.A.

## Author contributions

Conceptualization, V.D. and A.S.A.; Methodology, V.D., and A.S.A.; Investigation, V.D., and F.G.M.; Data curation, E.J.R.; Formal analysis, V.D., E.J.R., and S.A.M.; Writing – Original draft, V.D.; Writing – Reviewing and Editing, V.D., F.G.M., and A.S.A.; Visualization, V.D., F.G.M., and A.S.A.; Project administration, V.D. and A.S.A.; Funding Acquisition, A.S.A.; Supervision, A.S.A.

## Declaration of interests

The authors declare no competing interests.

## Supplementary figure legends

**Figure S1, related to Figure 1:**
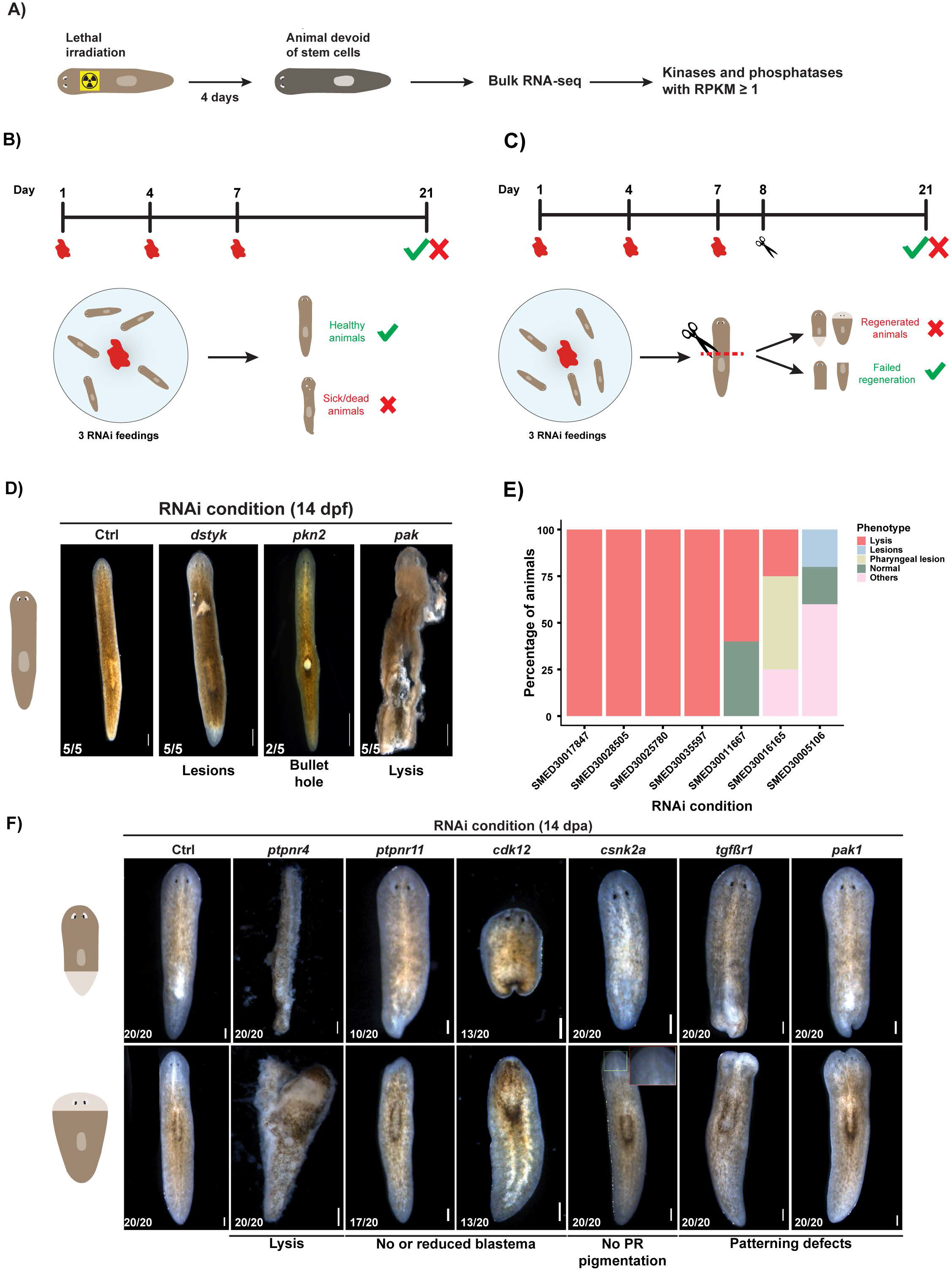
Two-part RNAi screen for kinases and phosphatases identified genes important for patterning. **(A)** Selection of kinases and phosphatases for RNAi screen. **(B)** RNAi feeding regimen and phenotype scoring criteria for homeostasis RNAi screen. **(C)** Cartoon representation showing RNAi feed schedule and scoring for the regeneration RNAi screen. **(D)** Live animal images at 14 dpf representing different phenotypes observed in the homeostasis screen. **(E)** Quantification of phenotypes observed in unamputated animals at 14 dpf. **(F)** Phenotypes identified in the regeneration screen showing posterior regeneration (top row) and anterior regeneration (bottom row). Scale bar: 200 um.

**Figure S2:**
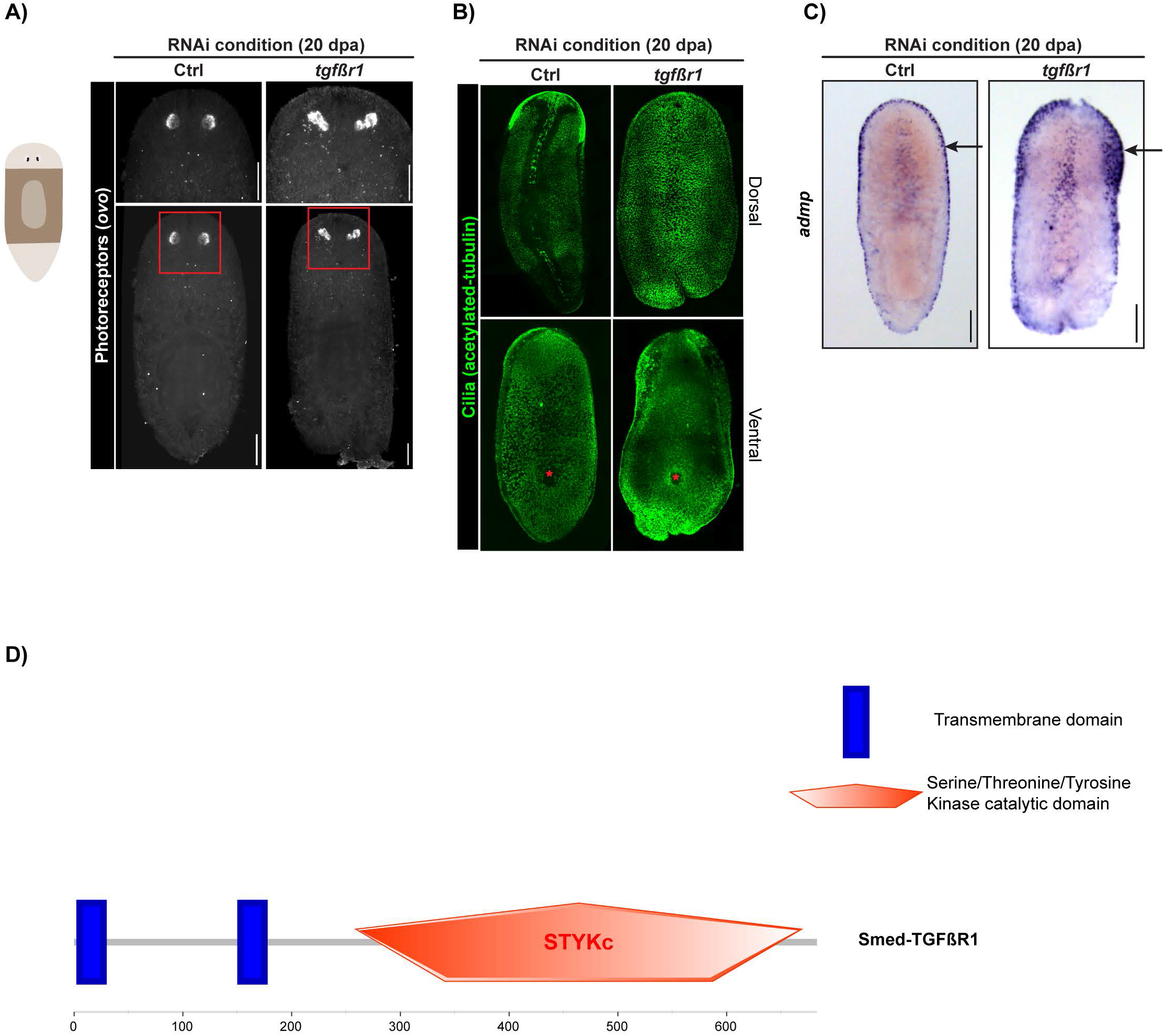
t*g*fβr1 is required for patterning the DV axis. **(A)** Maximum intensity projection images showing photoreceptors in regenerated trunk fragments at 20 dpa. **(B)** Dorsal and ventral cilia pattern in regenerated trunk fragments at 20 dpa. Pharynx is indicated by red star **(C)** Expression of *admp* at 20 dpa. Arrows indicate lateral expression of *admp* **(D)** Domain architecture of Smed-TGFβR1 protein. Scale at the bottom shows the amino acid position in the protein. Scale bars in the images are 200 um.

**Figure S3, related to Figure 1:**
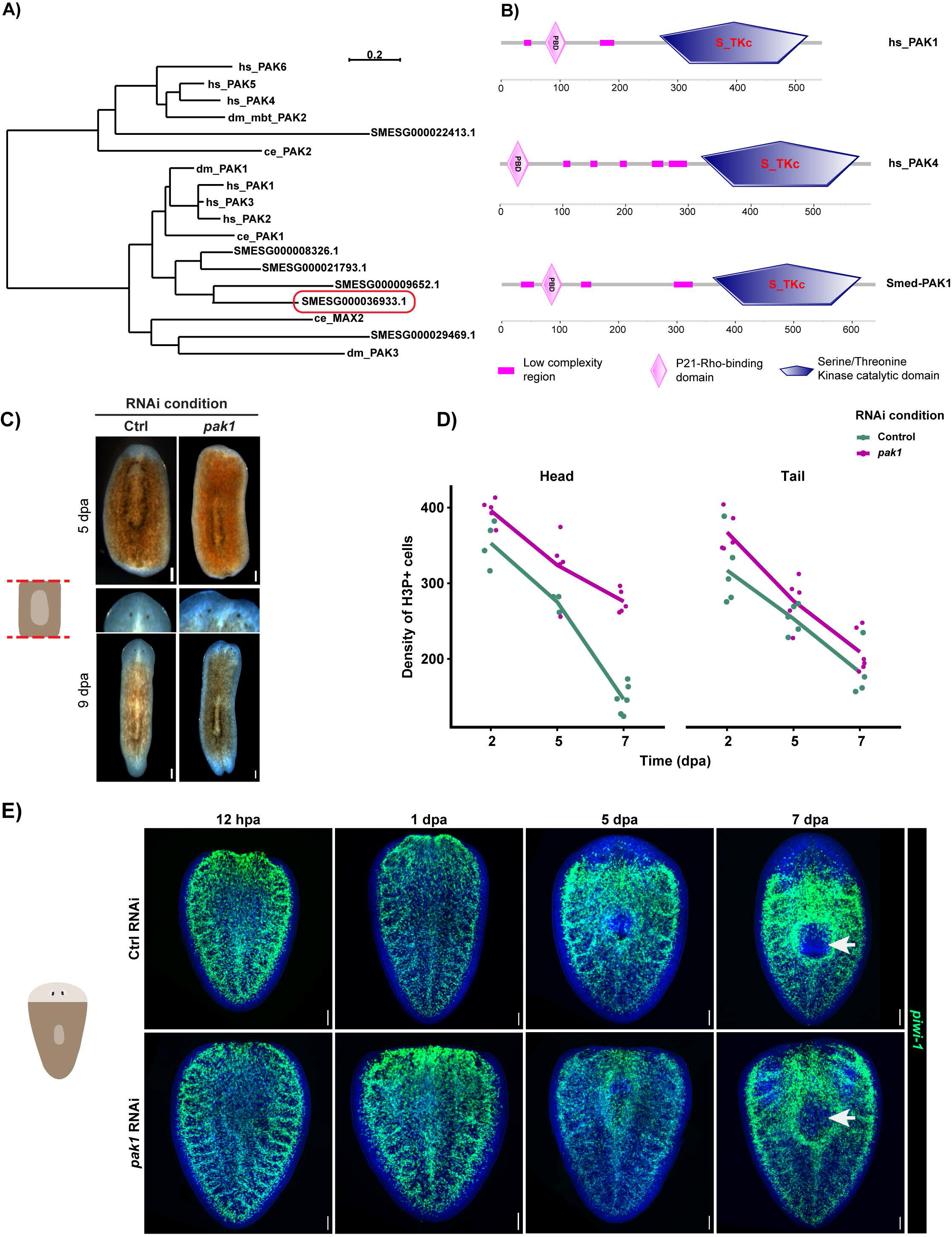
Smed-PAK1, a member of p21-activated family of kinases, regulates blastema formation without influencing amputation dependent stem cell responses. **(A)** Phylogenetic tree inferred from multiple sequence alignment of PAK kinases from *Homo sapiens* (hs), *Drosophila melanogaster* (dm), *Caenorhabditis elegans* (ce) and planarian *S.mediterranea* (SMESG). Smed-PAK1 from this study is highlighted with a red box. **(B)** Domains in human PAK1, PAK4 and Smed-PAK1 showing the N-terminal P21-Rho-binding domain (PBD) and the C-terminal Serine/Threonine Kinase catalytic domain (S_TKc). Scale below the domain structures indicate amino acid positions in the protein. **(C)** Images of regenerating trunk fragments at 5 and 9 dpa. **(D)** Densities of mitotic neoblasts during both posterior and anterior regeneration at 2, 5, and 7 dpa. **(E)** Maximum intensity images of FISH showing distribution of stem cells during anterior regeneration in tail fragments. White arrows indicate a region devoid of stem cells that regenerates a pharynx. Scale bar: 200 um.

**Figure S4, related to Figure 1 and 2:**
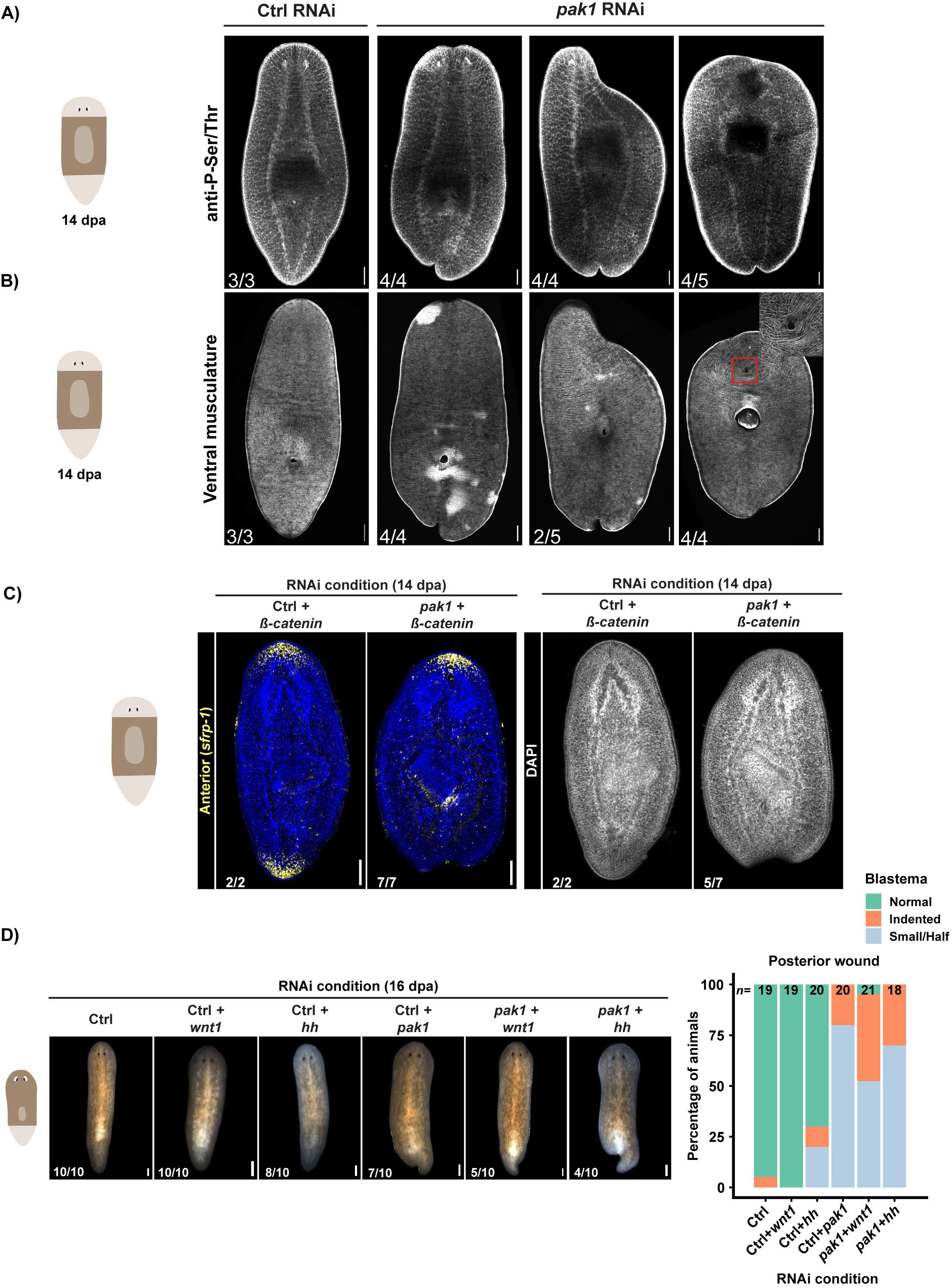
*pak1* modulates WNT/β-catenin signaling to pattern the AP axis. **(A)** Maximum intensity projections of immunostainings showing the cephalic ganglia, ventral nerve cods and photoreceptors in 14 dpa trunk fragments. **(B)** Maximum intensity projections of ventral third of the animals showing body wall musculature marked by 6GC10 antibody at 14 dpa. Area marked by red box is zoomed in and shows the formation of supernumerary mouth opening **(C)** Regenerated trunk fragments with expression of the anterior marker *sfrp-1* and DAPI staining in the same animals highlighting cephalic ganglia and ventral nerve cords. **(D)** Live animal images at 16 dpa showing posterior regeneration in double RNAi conditions and quantification of phenotypes in the plot on the right. Scale bar: 200 um.

**Figure S5, related to Figure 3 and 4:**
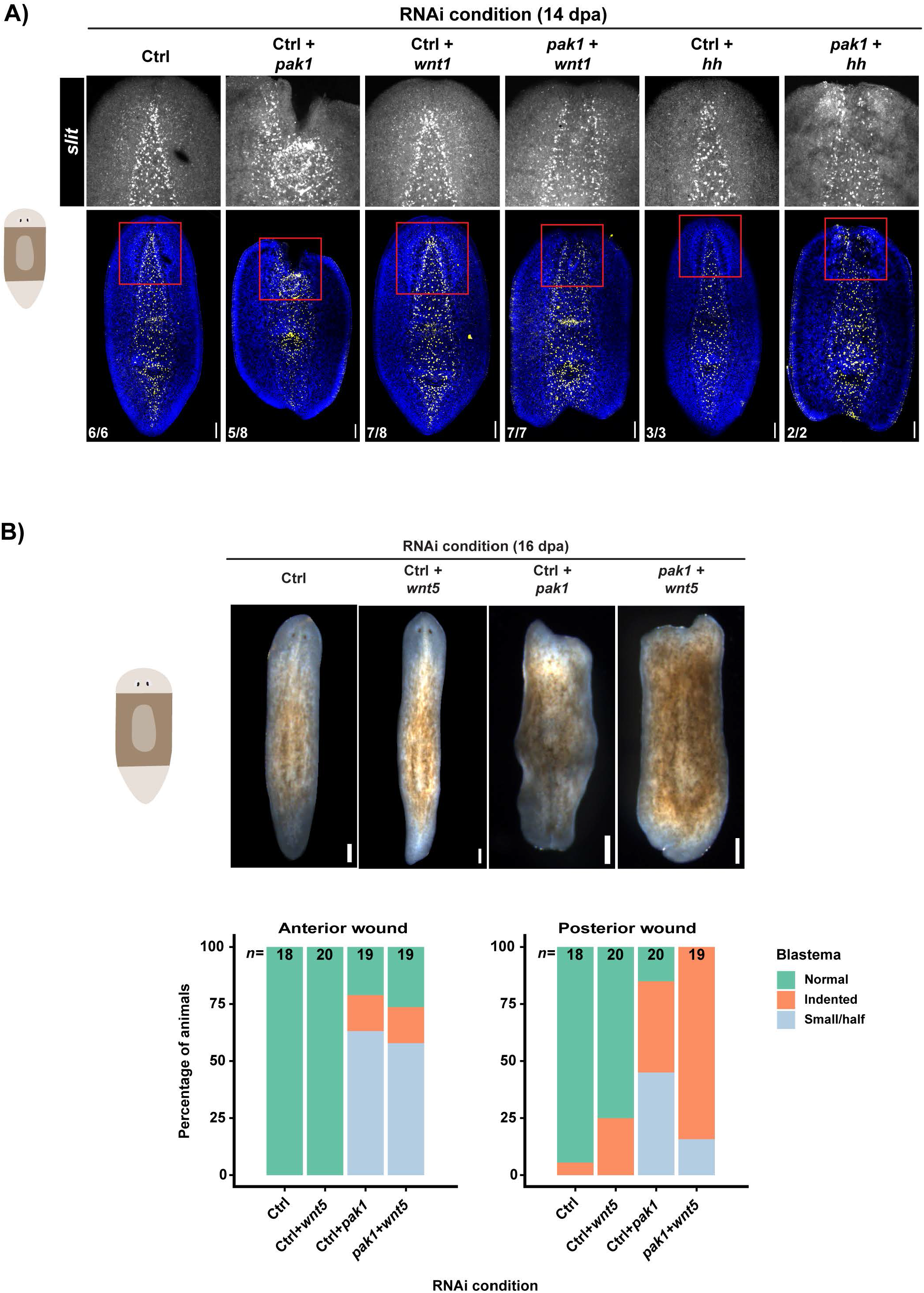
ML axis patterning by *pak1* is independent of WNT/β-catenin signaling and is synergistic with *wnt5*. **(A)** Maximum intensity projections of the ventral third of the animals showing expression of *slit* in 14 dpa trunk fragments. Areas marked by red are zoomed in the top row. **(B)** Live animal images of 16 dpa trunk fragments and quantification of phenotypes at both anterior and posterior wounds. Scale bar: 200 um.

**Figure S6, related to Figure 5:**
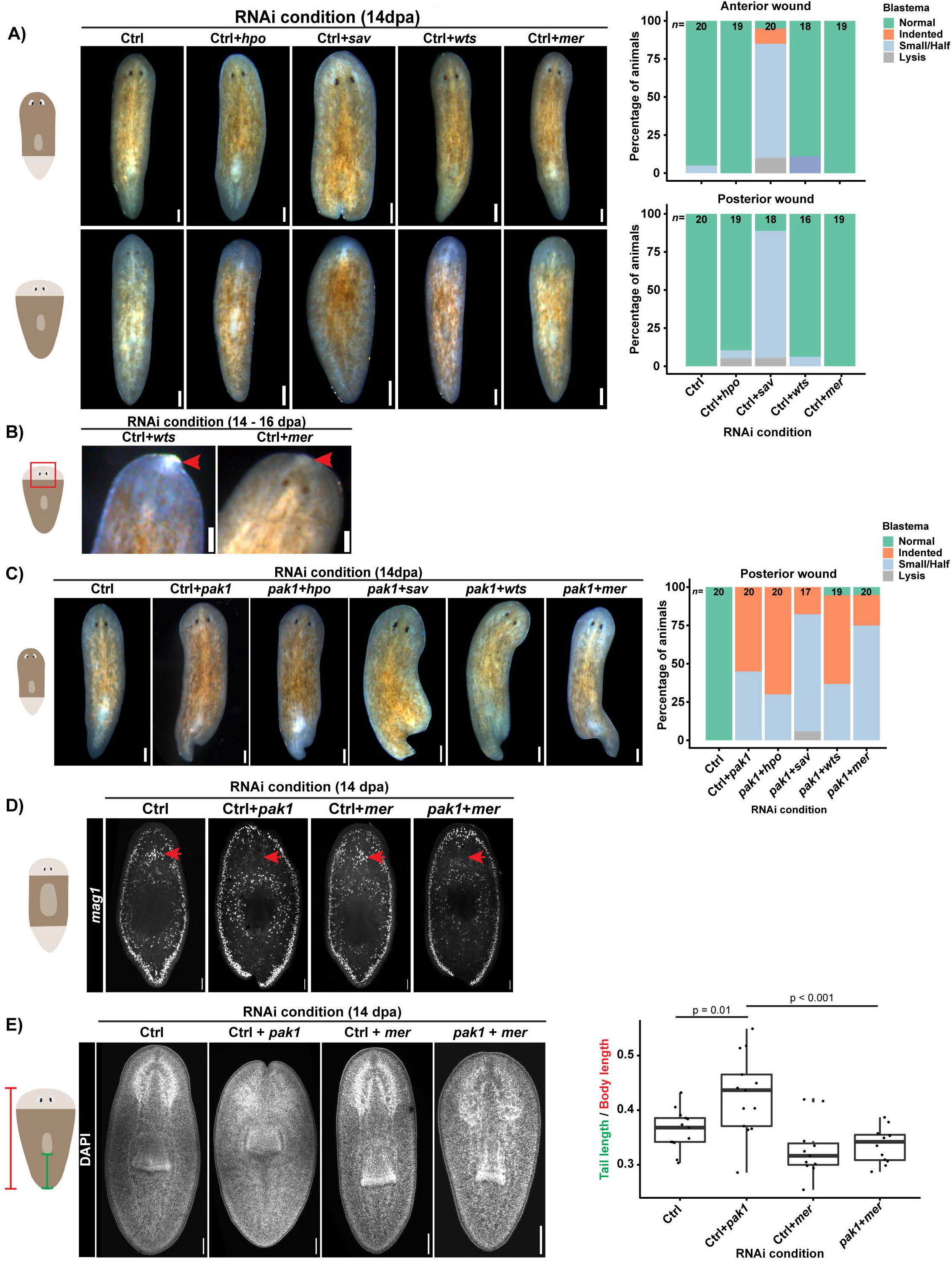
*mer(RNAi)* restores appropriate body proportions during anterior regeneration in *pak1(RNAi)* animals, but fails to rescue posterior regeneration. **(A)** Live animal images showing posterior regeneration (top row) and anterior regeneration (bottom row) at 14 dpa. Frequencies of regeneration phenotypes observed in the RNAi conditions are shown in the plots on the right side. **(B)** Live animal images of tail fragments at 14-16 dpa showing occurrences of rare phenotypes (red arrow). **(C)** Live animal images at 14 dpa of head fragments regenerating tails. Regeneration phenotypes observed in the double RNAi conditions are quantified in the plot to the right. **(D)** Maximum intensity projection images showing secretory cells marked by *mag1* in 14 dpa trunk fragments. **(E)** Maximum intensity projections of DAPI stained tail fragments at 14 dpa. Plot showing the ratio of tail length to body length. p-value is calculated from two-tailed Student’s t-Test. Scale bar: 200 um.

**Supplementary Table S1: Predicted kinases and phosphatases with primers used for cloning into RNAi vector.** Hits from homeostasis screen are highlighted in yellow and hits from regeneration screen are highlighted in green.

## Experimental procedures

### Animal husbandry

Asexual planarians of *Schmidtea mediterranea* (strain CIW4) were grown in 1X Montjuic salt in recirculation systems (Arnold et al., 2019). Animals were fed beef liver chunks, 1-3 times a week. Animals drawn from recirculation systems were maintained in static cultures at 20°C and starved for at least a week before using for experiments (Newmark and Sánchez Alvarado, 2000).

### Annotation of kinases and phosphatases

Kinases in *S. mediterranea* gene models (Rozanski et al., 2019) were annotated with HMMER (3.2.1; default parameters; e-value cutoff 1e-5) (Eddy, 2011) using HHM databases from Kinomer (Miranda-Saavedra and Barton, 2007) and EKPD (Wang et al., 2014). Phosphatases were annotated with HMMER (3.2.1; default parameters; e-value cutoff 1e-3) using HMM from EKPD. p21 PFAM (Mistry et al., 2021) domains were identified using HMMER run through Interproscan (Jones et al., 2014). Best hits to *Homo sapiens*, *Mus musculus*, *Drosophila melanogaster*, *Danio rerio* and *Caenorhabditis elegans* were determined via BLAST (e-value .001) (Altschul et al., 1990; Camacho et al., 2009). RNAseq TPMs are from Cheng et al., 2018 (GSE80562) (Cheng et al., 2018)

### Phylogenetic tree for PAK kinases

Published PAK kinases from *H. sapiens*, *D. melanogaster,* and *C. elegans* were combined with the six *S. mediterranea* kinases containing both kinase and p21 domains. These sequences were aligned with MUSCLE (version: v3.8.31) (Edgar, 2004), trimmed with trimAl (version: v1.4.rev15 ; parameters: -automated1) (Capella-Gutiérrez et al., 2009) and then concatenated and cleaned with phyutility (version: v.2.2.6; parameters -clean .5) (Smith and Dunn, 2008). Phylogenetic tree was created using RAxML-NG (version: 1.0.2; parameters: --model PROTGTR+G --seed 12345) (Kozlov et al., 2019).

### Cloning genes into RNAi vector

PCR Primers were designed via Primer3 (https://primer3.ut.ee/) to amplify transcript sequences from the Sánchez Alvarado lab transcriptome (Reference?). Amplicons were designed to be 400-600 bp in length. Flanking regions homologous to the pPR-T4P plasmid vector were added to the 5’ ends of the primers using which amplicons were inserted into the plasmids by Gibson assembly. Plasmids were transformed directly into *E. coli* strain HT115. Transformants were verified by sequencing. A list of primers for each kinase and phosphatase clones is included in Table S1. Primers for other clones used in the double RNAi experiments are provided in the table below. Gene sequence information is available at www.planosphere.stowers.org.

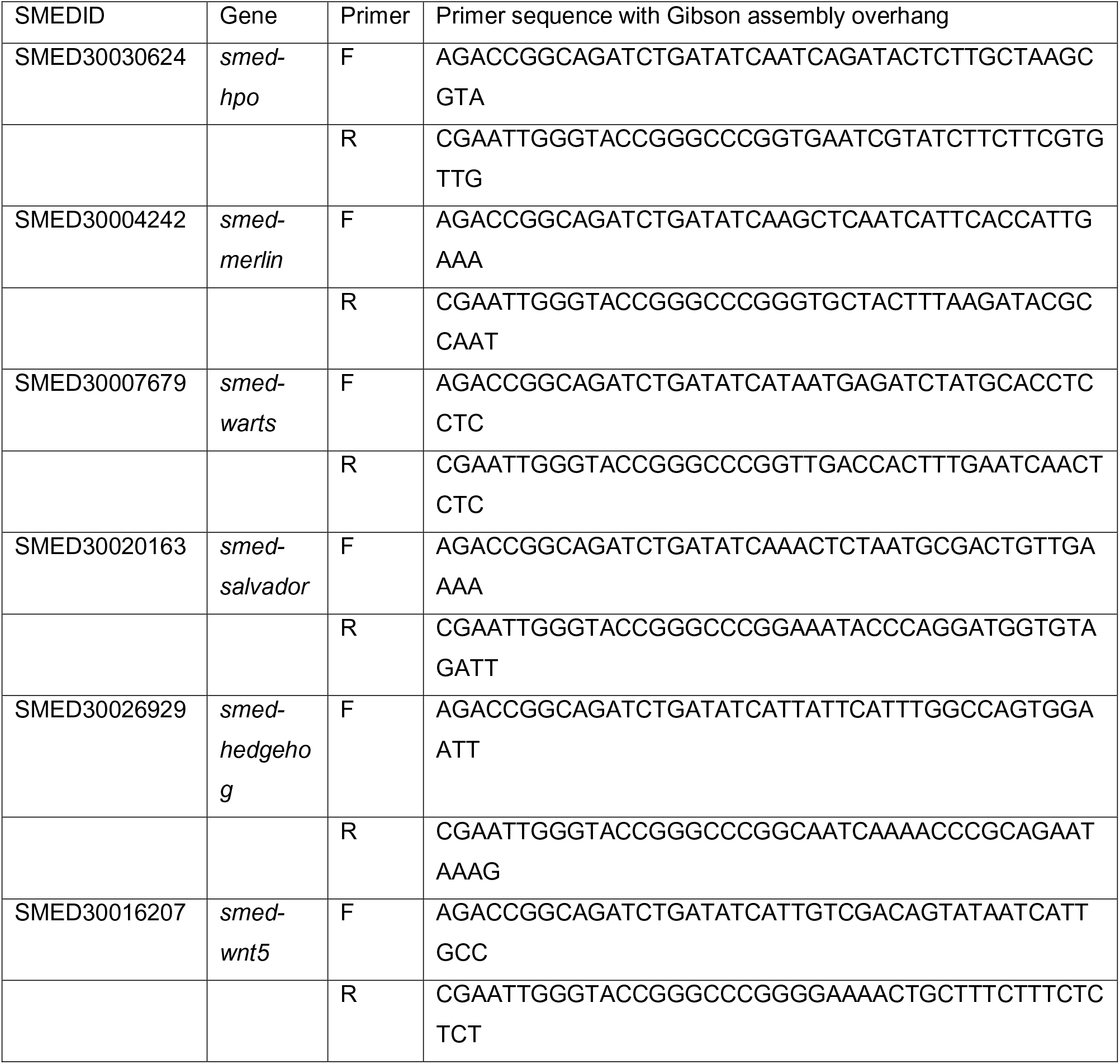

### High throughput RNAi food preparation and screening

Bacterial cultures were grown in 2X YT media supplemented with 50 ug/ml Kanamycin and 10 ug/ml Tetracycline in deep (10 ml), round bottomed 24-well plates. Initial culture of 2ml was grown for 16-18 hours at 37 °C. Double stranded RNA production was induced by diluting the overnight culture with 6 ml of 2X YT with 50 ug/ml Kanamycin, 10 ug/ml Tetracycline and 1 mM IPTG. After 4 hours of induction bacterial cultures were pelleted at 1000 rpm for 10 minutes. Each bacterial pellet was mixed with 62.5 ul of homogenized beef liver mixed with food coloring in 1X Montjuic salts. Bacterial pellets and homogenized beef liver were mixed by vortexing for 10 minutes. *C.elegans* gene *unc-22* with no nucleotide homology to any planarian sequence was used as a control RNAi. RNAi food was stored at −20C in Ziploc bags during the duration of RNAi feedings.

Animals were housed in cups with meshes in a flow through system for the duration of the screen (Arnold et al., 2016). Each RNAi condition had ∼5 animals and they were fed 15 ul of RNAi food for 3 times in a span of 7 days. For the regeneration screen animals were amputated pre-pharyngeally a day after the last feeding. Animals were observed for phenotypes for the next two weeks and were scored on either 14 days post feeding (dpf) or 14 days post amputation(dpa).

### RNAi

RNAi in planarians was induced by feeding animals with bacteria expressing dsRNA using the “soft serve” method as previously described (Gurley et al., 2008; Rink et al., 2009). An overnight bacterial culture was diluted to 10% in 2X YT media complemented with 50 ug/ml Kanamycin and 10 ug/ml Tetracycline. After 4 hours of IPTG induction, cultures were spun down at 4000 rpm for 10 minutes. Bacterial pellet corresponding to 50 ml of culture was mixed with 150 ul of homogenized beef liver with food coloring. When the bacterial cultures were mixed to make double RNAi food, the ratio of bacteria to liver was doubled so that the concentration of each gene was the same as in the single knockdown. In all experiments animals were fed 6 times except the following – in Figure S1D, S1F, 2B, 3C, and 3D animals were fed 3 times and in Figure S4C animals were fed 5 times with Control or *pak1(RNAi)* food and once with β*-catenin(RNAi)* or *pak1+* β*-catenin* RNAi food.

### *in situ* hybridization and immunofluorescence

Whole mount in situ hybridizations were performed as mentioned previously with either NAC treatment (King and Newmark, 2013; Pearson et al., 2009) or nitric acid/ formic acid (NAFA) protocol (Guerrero-Hernández et al., 2021).

When using the NAC treatment, animals were euthanized with 5% NAC for 5 minutes and then fixed with 4% formaldehyde for 45 minutes. To increase the permeability of the tissue, samples were treated with the reduction solution for 10 minutes. Samples were then washed with PBS + 0.3% Triton-X, following which were dehydrated in methanol and stored in −20 °C for at least an hour. When ready, samples were rehydrated in PBS + 0.3% Triton-X and bleached using Ryan King’s formamide bleach solution for 2 hours (King and Newmark, 2013). After bleaching, animals were treated with Proteinase K (2 ug/ml) for 10 minutes to promote probe permeabilization.

With NAFA protocol, animals were euthanized using nitric acid and fixed for 45 minutes with 4% paraformaldehyde in the presence of 4.8% formic acid. Samples were dehydrated in methanol and stored at −20 °C until ready to process for *in situ* hybridization. Animals were rehydrated in PBS + 0.3% Triton-X and bleached for 2 hours in 6% H_2_O_2_ and 1% formamide.

Probe hybridization and signal development for the samples fixed by either NAC treatment or NAFA protocol were the same and are described briefly. Hapten labeled antisense probes were hybridized for >16 hours at 56 °C. After a series of washes with Wash Hyb and SSCx + 0.1% Tween-20, samples were blocked with 5% horse serum + 0.5% Roche western blocking reagent for 2 hours. Following blocking, samples were incubated overnight with appropriate antibody. Post washes with MABT, signal was developed with either NBT-BCIP (colorimetric) or fluorescently tagged tyramides. Animals were optically cleared in Scale A2 + DABCO before imaging (Lei et al., 2016).

For immunostaining assays animals were fixed and bleached as mentioned in the NAFA protocol (Guerrero-Hernández et al., 2021). After bleaching samples were blocked with 5% goat serum or 10% horse serum in PBS + 0.3% Triton-X. Blocked samples were incubated with appropriate primary antibodies for overnight at room temperature. The samples were washed 6 times for 20 minutes each with PBS + 0.3% Triton-X. Following washes, samples were incubated overnight at room temperature with fluorescently tagged secondary antibodies and DAPI. Samples were cleared in Scale A2 + DABCO before imaging.

| Antibody | Supplier | Catalog No. | Dilution |
| --- | --- | --- | --- |
| Anti-DIG-AP | Roche | 11093274910 | 1:3000 |
| Anti-DIG-POD | Roche | 11207733910 | 1:1000 |
| Anti-Fluorescein-POD | Jackson Laboratories | AB_2314402 | 1:2000 |
| Anti-H3P | Abcam | ab32107 | 1:500 |
| Anti-P-S/T | Cell Signaling | #9631 | 1:200 |
| 6GC10 (muscle) | DSHB | 6G10-2C7 | 1:400 |
| Anti-Synorf | DSHB | 3C-11 | 1:100 |
| Anti-arrestin | Lagen Labs | 016-arrestin-01 | 1:200 |
| VC-1 | Kind gift from K. Agata and D. Palakodeti | NA | 1:2500 |
| Anti-acetylated tubulin | Cell Signaling | #5335 | 1:500 |
| Anti-mouse-488 | Abcam | 150113 | 1:500 |
| Anti-mouse-555 | Abcam | 150118 | 1:500 |
| Anti-rabbit-488 | Abcam | 150073 | 1:500 |
| Anti-rabbit-555 | Abcam | 150086 | 1:500 |
| Anti-rabbit-647 | Abcam | 150083 | 1:500 |

### Imaging and image analysis

Live animal images and images from colorimetric WISH samples were obtained on Leica M205 microscope. Fluorescent images were taken on a Nikon Ti Eclipse equipped with a Yokogawa CSU W1 spinning disk head and Prior plate loading robot. Slides were loaded automatically, and animals found using custom software as in (Adler et al., 2014). Stitching and processing were performed in Fiji (Schindelin et al., 2012) using plugins from the Stowers Fiji update site as well as macros at https://github.com/jouyun/2021_Doddihal. Images were processed in Fiji. For generating images showing the ventral third of the animals, images were sub-stacked into three equal parts and Z-slices corresponding to the ventral side of the animal were used to generate maximum intensity projections.

## Notes

### Competing Interest Statement

The authors have declared no competing interest.

